# Gels for Live Analysis of Compartmentalized Environments (GLAnCE): A Tissue Model to Probe Tumour Phenotypes at Tumour-Stroma Interfaces

**DOI:** 10.1101/782086

**Authors:** Elisa D’Arcangelo, Nila C. Wu, Tianhao Chen, Andi Shahaj, Jose L. Cadavid, Linwen Huang, Laurie Ailles, Alison P. McGuigan

**Affiliations:** Insitute of Biomaterials and Biomedical Engineering, University of Toronto, Toronto, CA; Department of Chemical Engineering and Applied Chemistry, University of Toronto, Toronto, CA; Princess Margaret Cancer Centre, University Health Network, Canada

**Keywords:** tumour-stroma interface, invasion, CAF, invasion-permissive matrix, EMT

## Abstract

The interface between a tumour and the adjacent stroma is a site of great importance for tumour development. At this site, carcinoma cells are highly proliferative, undergo invasive phenotypic changes, and directly interact with surrounding stromal cells, such as cancer-associated fibroblasts (CAFs) which further exert pro-tumorigenic effects. Here we describe the development of GLAnCE (Gels for Live Analysis of Compartmentalized Environments), an easy-to-use hydrogel-culture platform for investigating CAF-tumour cell interaction dynamics *in vitro* at a tumour-stroma interface. GLAnCE enables observation of CAF-mediated enhancement of both tumour cell proliferation and invasion at the tumour-stroma interface in real time, as well as stratification between phenotypes at the interface versus in the bulk tumour tissue compartment. We found that CAF presence resulted in the establishment of an invasion-permissive, interface-specific matrix environment, that leads to carcinoma cell movement outwards from the tumour edge and tumour cell invasion. Furthermore, the spatial stratification capability of GLAnCE was leveraged to discern differences between tumour cell epithelial-to-mesenchymal (EMT) transition genes induced by paracrine signaling from CAFs versus genes induced by interface-specific, CAF-mediated microenvironment. GLAnCE combines high usability and tissue complexity, to provide a powerful *in vitro* platform to probe mechanisms of tumour cell movement specific to the microenvironment at the tumour-stroma interface.

## Introduction

Stromal tissue compartments actively maintain epithelial proliferation and differentiation in adult organs, by modulating the extracellular matrix architecture of the tissue niche (1–4). The compartmentalization between stromal and epithelial tissue units, present during homeostasis, is lost in neoplasia, where the compartment-separating basement membrane is broken down (5–7). This allows contact between epithelial and stromal cell populations and interspersing of these populations at the tumour invasive front (8–10). Stromal fibroblasts become activated and acquire a cancer-associated phenotype as a result of, or concurrently with, carcinoma cell movement into the stromal compartment, (8,11,12). These cancer-associated fibroblasts (CAFs) are often the most abundant stromal cell type (13) and readily infiltrate carcinomas, beginning at the tumour-stroma interface. At the tumour-stroma interface, CAFs facilitate increased tumour aggressiveness by increasing tumour cell proliferation and dissemination of invasive cell clusters (14)(15–17), both directly through secretion of various growth factors and cytokines (11,18,19) and indirectly, by promoting angiogenesis (20). Further, CAFs directly initiate tumour cell invasion by supporting epithelial-to-mesenchymal transitions (21–23) and by altering the composition and mechanical properties of tumour extracellular matrix (ECM) (24–27). As a result, patients with tumours characterized by high stromal content carry a worse prognosis in a multitude of solid tumours (28–31).

While there is ample evidence showing that CAFs are major players in the tumour microenvironment (TME), and that specific invasive tumour cell phenotypes occur at the tumour-stroma interface (32,33), there is a lack of *in vitro* models capable of stratifying CAF-tumour cell interactions in the context of a relevant tumour tissue architecture. Such a platform would enable analysis of the tissue location-specific contributions of CAFs on tumour cell phenotype and enhance our understanding of the spatial and temporal dynamics at the tumour margin, compared to the bulk tumour tissue, which may uncover novel treatment avenues. Typical *in vitro* invasion assays involve culture of tumour cells in a hydrogel matrix, where invasion into the hydrogel phase is monitored either in real time or after fixation. Cocultures of multiple cell types can be incorporated into the hydrogel phase, in order to attain more sophisticated cellular environments (34–36). Experimental set ups span a range of complexities from simple hydrogel plugs in a multi-well plate (37) and filter insert (38), to more sophisticated microfluidic platforms where different cell populations can be patterned into specific structures and tissue compartments. These microfluidic approaches have enabled real time monitoring of tumour cell invasion into neighbouring stromal tissue (39–41) and revealed dynamic events such as the transition to an invasive phenotype, colonization of adjacent compartments, and tumour cell extravasation. These changes in cell migration, morphology, and proliferation would be particularly challenging to quantify in animal or other *in vitro* culture models.

While microfluidic platforms (39,42) offer a powerful approach for understanding differences in CAF-tumour interactions at the interface versus in the bulk tissue, the often sophisticated setups of these platforms come at the price of usability: microfluidic devices are often cumbersome to assemble, require external pumps for prolonged culture times, and are incompatible with existing work flows and instrumentation (43). Furthermore, cell densities in closed culture channels are often limited to avoid detachment of the hydrogel-cell phase from the channel walls and hypoxic cores within the cultures. Overall, this has led to poor adoption of microfluidic devices for cell culture among the wider biology research community, where 2D cultures (44), aggregate cultures (45) and simpler hydrogel plug/dome culture arrangements (43) are predominant.

In this report, we describe the development of GLAnCE (Gel for Live Analysis of Compartmentalized Environments), an easy-to-use culture platform that recapitulates the overall architecture of the invasive tumour front. Uniquely, this platform allows real-time visualization of the interaction between cell populations within the context of an architecturally relevant tissue geometry. Using GLAnCE, we characterized tumour cell movement at the tumour-stroma interface and its dependency on CAF interactions. We show that squamous cell carcinoma (SCC) cells at the invasive front require CAFs for the establishment of an invasion-permissive TME, leading to enhanced growth and invasion. Finally, we demonstrate the use of GLAnCE for identifying tumour – stroma signalling that promotes EMT specifically at the invasive front.

## Materials and Methods

### Cell culture

Human squamous cell carcinoma cells of the tongue (CAL33 cells) were a gift from A. Nichols from London, ON, and were stably transfected with an mCherry construct (Clontech, Mountain View, CA, USA), using lentivral transduction as described previously (46). MDCK cells were purchased from ATCC. Three CAF lines (ID numbers 61137, 65055, 61162) were isolated from three different patient samples in compliance with the University Health Network Ethics Board guidelines and CAF identity was confirmed for each line as CD45-, CD31-, cytokeratin- and vimentin+, as described before (46). CAFs were further STR profiled to ensure a match with the donor biopsy tissue. CAFs were tagged with eGFP by lentiviral transduction (46). MDCK cells were grown in Eagle’s MEM (Millipore Sigma, USA) supplemented with 10% FBS (12483020, Gibco, CA) and 1% penicillin/ streptomycin solution (30-002-CI, Corning, USA). CAL33 tumour cells and CAFs were maintained in IMDM (Gibco, USA), supplemented with 10% FBS and 1% penicillin/ streptomycin solution, in tissue culture polystyrene flasks prior to use for compartmentalized cultures. CAFs were used up to and including their 10th passage. Cultures were maintained at 37 °C in 5% CO2.

### Fabrication of device components

An array of 4×6 channel-shaped features of 276 μm +/- 31 μm was micro-milled into the surface of an aluminium slab (6061-T6, McMaster-Carr, CA) of 117 cm x 78 cm and fine grit sand-paper (grit size 2500, McMaster-Carr, CA) was used to polish the features’ surface. This piece was subsequently utilized as the stamp for producing GLAnCE polystyrene (PS) device bottoms. Specifically, the stamp was pushed into a sheet of virgin crystal PS (ST313120, GoodFellow, UK) with 1700N of force and at 190 °C for 6 minutes, with a temperature ramp and de-ramp of 50 °C /minute. After PS de-embossing from the stamp, the PS device bottoms were trimmed to fit the overall dimensions of a bottom-less 24 well plate (662000-06, Greiner Bio-One, Austria). The channel-shaped depression in the PS were termed ‘open channels’. PS device bottoms were exposed face-up to UV light at 254 nm and 15 Watts for 150 min. This step increased hydrogel adhesion to the PS surface, by breaking the aromatic rings (photooxidation of PS by carbon-carbon bond scission) exposed on the PS surface, which gives rise to -CH3 groups (47); it has been observed that this process increases the adhesiveness of PS towards collagen solutions by allowing more efficient surface entanglement of collagen fibrils (48).

Polydimethylsiloxane (PDMS) slabs of approximately 0.5 cm thickness were generated by pouring liquid PDMS, at a base to crosslinker ratio of 10:1 (4019862, Sylgard 184, Ellsworth Adhesives, USA) into a 9 cm x 13 cm, rectangular plastic mold, which was baked at 68 °C for a minimum of 2 hours. Upon solidification, PDMS slabs were removed from the mold inside a clean room, aligned with a template PS device bottom, and 4×24 through-holes were manually punched through the PDMS using a 1 mm-bio punch (Harris Uni-Core, Ted Pella, USA). This yielded 2 hydrogel inlet and 2 air outlet holes for each open channel on the template PS device bottom. PDMS slabs were wrapped in dust-capturing tape (7922A5, McMaster-Carr, CA) for storage. A sheet of double-sided, poly-acrylic adhesive tape (ARcare^®^ 90106, Adhesive Research, USA) was cut to the overall dimensions of 117 cm x 78 cm using a Silhouette Cameo Electronic Cutting Machine (Silhouette, USA) to remove circular areas of tape corresponding to the wells of the no-bottom 24 well plate. All parts of the device, including the PS device bottom, PDMS slab, bottom-less 24 well plate and adhesive tape were UV-sterilized for 15min on each side prior to assembly and use. PDMS slabs and PS device bottoms were sandwiched together and incubated at 37 °C until the time of hydrogel injection, in order to allow for temperature-induced expansion of the materials to occur. Sandwiching of the PS channels against the PDMS resulted in temporarily ‘closed’ channel configurations.

### Sample preparation for hydrogel culture

Rat tail collagen type I at 5mg/ml was prepared according to manufacturer’s protocol (50201, iBiDi, Germany) to obtain a collagen solution of 3mg/ml. Rat tail collagen type I at 10mg/ml (50205, iBiDi, Germany) was neutralized according to the same protocol. Bovine collagen type I at 3mg/ml (5005-B, Advanced BioMatrix, USA) was prepared according to manufacturers’ instructions. GFR Matrigel (356231, Corning, USA) was utilized undiluted from stock. All hydrogels were kept on ice until use and, for neutralized collagen solutions, for no longer than 10 min.

### Generation of compartmentalized hydrogels and GLAnCE culture device

Following a standard trypsinization protocol, cells were spun down and resuspended in the relevant hydrogel at a cell density of 1.47 × 10^6^ cells/ml for tumour cells and 2.94 × 10^6^ cells/ml for CAFs and placed on ice. Subsequently, 3.4 μl of hydrogel cell suspension, containing 5000 tumour cells, was injected into the right closed channel inlet holes present in the PDMS-PS device bottom sandwich using a standard pipette with tip diameter of 1mm. This was carried out on a foam pad heated to 37 °C, which helped minimize the material contraction at room temperature, as well as ensuring initiation of hydrogel gelation at the correct temperature. The PDMS-PS sandwich was subsequently placed at 37 °C for 5 min and covered with a sterile, PBS-soaked Kim Wipe to prevent hydrogel evaporation. After 5 min, a second 3.4 μl - volume of hydrogel-cell solution, containing 10,000 CAFs, was injected into the closed channels via the right inlet holes and the device was placed, covered by a moistened Kim Wipe, at 37 °C for 20 min. Finally, the PDMS slab was peeled off using tweezers, exposing the array of 24 compartmentalized hydrogels, supported within the PS device bottom, which was attached to a no-bottom, 24 well plate via a polyacrylic adhesive tape, resulting in a fully assembled, 24 well culture plate. Media was added to each well for culture.

Cancer cell lines are typically highly proliferative, especially when grown in high serum conditions (e.g. 10% FBS). Accordingly, substantial hydrogel crowding was detected in compartmentalized culture above a certain seeding density of tumour cells, which interfered with the identification of invasive strands in fluorescent micrographs and drastically increased the outwards movement of the tumour compartment at the interface. Specifically, tumour cells formed large 2D sheets at the bottom of the hydrogel culture and adhered to the underlying PS substrate. A critical cell number of tumour cells was therefore selected for all experiments, to avoid significant numbers of cells growing as 2D sheets to ensure that proliferation and invasion in 3D were the predominant mechanism of interface movement.

### Nuclear, Viability, MTT and EF5 staining

Live nuclei were stained with Hoechst (1:10,000) and washed in PBS prior to confocal image acquisition. Viability was assessed by incubating samples with 0.5μl/ml calcein AM for live cells and 2μl/ml ethidium homodimer-1 to detect dead cells (L3224, Thermo Fisher Scientific, USA) for 30 min, followed by media replacement and quantification of % live cells in Fiji (49).

Metabolic activity was quantified by applying 20 μl of MTT (3-(4,5-Dimethylthiazol-2-yl)-2,5-Diphenyltetrazolium Bromide, 5 mg/ml, M6494 Thermo Fisher Scientific, USA) to wells on day 5 of culture and samples were incubated in the dark at 37 °C for 2 h. The samples were then imaged in brightfield mode and MTT intensity quantified in Fiji as mean grey value. The presence of oxygen gradients/hypoxia was investigated using EF5 staining as described previously (46,50). In brief, 100 mM of EF5 was applied for 3h to compartmentalized hydrogels laden with tumour cell aggregates at day 5 of culture. Following fixation in 4% PFA, samples were permeabilized with 2.0% Triton-X, blocked in 2.5% BSA, and stained with Cy3-conjugated anti-EF5 antibody (1:150, ELK3-51), as well as DRAQ5 (1:1000) (Danvers, MA). This was carried out for hydrogels in regular culture at 5% CO2, hydrogels cultured in anoxia, using Hypoxystation (Don Whitley Scientific, UK) (in which case the EF5 was added 3hrs prior to the end of the 6hr anoxic incubation time), which served as a positive EF5 control, and normoxic cultures without the addition of the EF5 drug, which served as the negative control. EF5 images were obtained with confocal imaging and intensity levels were quantified as the mean grey value of the Cy3-conjugated ELK-351 antibody signal (as described at http://hypoxia-imaging.org/v2/methods/ef5manual.htm) in regions corresponding to cell nuclei, as defined by DRAQ5 positivity using Fiji.

### Confocal and widefield fluorescence microscopy

All confocal images were acquired on live samples using a Carl Zeiss LSM700 and 10 X magnification. For hypoxia quantification, 2 μm thick optical slices were imaged on a Leica SP8 at 20 X magnification; the gain was first set based on the maximum binding positive control and EF5 values were normalized to the range between negative and positive controls. All widefield images were obtained at 4 x magnification, using an ImageXpress Micro (Molecular Devices, USA) for high-throughput and time-lapse imaging. For the comparison between non-sliced and multi-slice + deconvoluted images, the focal plane was set at z-positions incrementally distant from the device bottom (z-slice thickness of 10μm) and 20 images were acquired per sample. These were subsequently deconvoluted using the 2D No Neighbor deconvolution algorithm in MetaXpress^®^ (Molecular Devices, USA).

### Staining Interconnected Hydrogel Compartments

Prior to seeding hydrogels, 3 mg/mL rat tail collagen type 1 was mixed 1:10 with AF647 (A21463, Invitrogen, US) or AF546 (A21123, Life Technologies, US) secondary fluorescent antibodies and compartmentalized hydrogels were then seeded as previously described. Confocal imaging was performed on the interface region.

### Automated quantification of compartment interface movement

Widefield fluorescent images were uploaded to MATLAB (Mathworks), where both, day 0 and day 5 images from the same sample were converted to binary images. In order to trace interface lines between non-contiguous aggregates, a number of image dilation and erosion steps were carried out and images were subsequently cropped to channel size. Using the MATLAB function bwtraceboundary (51), a line between non-zero pixels (i.e. cell fluorescence) and zero pixels (i.e. background) in both day 0 and day 5 images was traced, and the coordinates recorded. Then, the traced day 0 and day 5 images were overlaid and the area between the traces was isolated as a binary image with the function poly2mask (51). The area between traces was calculated by counting the number of pixels in the region. Finally, to calculate distance of cell movement and area occupied by cells, the number of white (cell-occupied) pixels in each pixel row along the x-axis of the image was quantified.

### Quantification of strand structure morphology and number of strand structures

Widefield fluorescent images of invasive tumour aggregate morphologies were obtained at 10 x magnification. Aggregate perimeters (P_aggregate_)were manually traced in Fiji and perimeter length was recorded. Using these values, corresponding theoretical circle area values were calculated and their theoretical circle perimeters (P_theoretical_) noted. By computing P_aggregate_ / P_theoretical_, a deviation from circularity was calculated, which increased with increasing aggregate elongation and approached 1 for near-circular aggregates. The number of strand structures in any given hydrogel was determined manually in Fiji, first by subtracting a PS channel background image (i.e. a PS channel filled with media only) from the micrograph to be quantified and then identifying strand structures based on their elongated morphology.

### Inhibition of tumour cell proliferation using mitomycin C (MMC)

Tumour cell proliferation was blocked by treating cells in 2D cultures with 10 µM MMC for 2 hours prior to generating GLAnCE compartmentalized cocultures.

### Conditioned media experiments

Conditioned media was harvested at day 5 from 3D hydrogel cultures and concentrated in 3 kDa-membrane concentration columns (Amicon^®^, MillioreSigma, USA) by spinning at 3200 x g for 60 min at 4 °C. Media concentrates were diluted in serum-free IMDM (+1% penicillin/ streptomycin) to half the total media volume originally collected, and topped up to the total original media volume with IMDM + 10% FBS +1% penicillin/ streptomycin. Conditioned media was stored at -20 °C and freeze-thawed a maximum of 3 times.

### Comparison of GLAnCE and dome cultures using fluorescent microspheres

A 96-well, flat-bottom tissue-culture plate and GLAnCE components were pre-warmed in a 37°C incubator. Uniformly dyed Dragon Green fluorescent (480, 520) carboxyl-functionalized polymer microspheres, with a mean diameter of 15.32 μm (Bangs Laboratories, Inc., USA), were re-suspended in 3 mg/mL of rat tail collagen. The collagen-microsphere mixes were prepared at a density of 10 000 microspheres/6.5 μL collagen and hydrogel droplets of equal volume (6.5μl) were deposited both as domes in the pre-warmed 96 well plate (1 dome/well) and as channel-shaped hydrogels in GLAnCE. After hydrogel gelation at 37°C for 30 minutes, PBS was added to each hydrogel and images were taken at 10X magnification in widefield fluorescence mode, using the FITC filter (emission excitation 495/519). The top surface of every GLAnCE hydrogel and dome was set as the z-plane of best focus. For image analysis, performed in Fiji, images were blurred with a median filter of radius 3 to reduce random noise. For the GLAnCE hydrogels, a rectangular ROI was drawn, in order to exclude hydrogel areas close to the channel inlets. For the gel domes, a circular region centered on the centroid of the image was drawn; the diameter of this region was determined manually for a randomly chosen image and applied to all dome images. Two classes of object areas were defined in micrographs: Firstly, the area of objects in focus (A_focus_) was identified based on object signal intensity, which exceeded a certain, manually set threshold value (kept constant across all images). Secondly, an area corresponding to the total number of objects (A_total_) was defined using a lower threshold value that included all objects present. The percentage of object area in focus was then calculated by computing A_infocus_/A_total_ *100.

### Assessment of tumour cell proliferation via EdU staining

To demonstrate the pro-proliferative effect of CAFs on CAL33 tumours cells at the interface, the Click-iT^®^EdU Cell Proliferation Imaging Kit with was used. At day 5 of culture, 300 uL of 10 μM EdU (Click-iT^®^EdU Cell Proliferation Imaging Kit) was added to compartmentalized hydrogels and returned to 37°C to incubate for 48 hours. Subsequently, each well was fixed with 4% PFA, permeabilized with 0.5% Triton X-100 and blocked with 3% BSA. Click-iT^®^ reaction cocktail, containing Alexa Fluor^®^ 488 azide (Molecular Probes^®^, USA), was added to each compartmentalized hydrogel for 30 minutes in the dark. Then, hydrogels were washed with 3% BSA. EdU images were obtained at and away from the interface with confocal imaging, with 2 μm thick optical slices. The percentage of proliferating cells was quantified using FIJI by first thresholding the EdU and mCherry fluorescent signals, then multiplying the thresholded images using FIJI’s image calculator process. The percentage area of the resulting image (EdU^+^ mCherry^+^ signal) was then divided by the percentage area of the mCherry signal.

### RT^2^ qPCR EMT Array

Compartmentalized Hydrogels were generated as compartmentalized monocultures, compartmentalized cocultures, and mixed cocultures, with a constant number of tumour cells/unit volume and ratio of tumour cells to CAFs. After 5 days, hydrogels were digested by incubation in a digestion buffer (collagenase, proteas, DNase, 50nM MgCl_2_ and Modified Earle’s Salt Solution (made in house)) for 45 min at 37 °C on a shaker at 600 rpm, which released all cells from the gels. Upon neutralization of the digestion buffer, cells were pelleted and resuspended in sorting buffer (5% FBS in PBS), stained for viability (DAPI, 1:1000), and mCherry^+^ tumour cells were isolated using fluorescence-activated cell sorting (FACS). A purity check on 1000 cells/sample was performed to confirm absence of GFP^+^ CAFs. The isolated tumour cell population was subsequently subjected to RNA extraction (RNeasy Mini Kit, 74104, Qiagen, Germany) and spectrophometric concentration measurements (NanoDrop^TM^, Thermo Fisher Scientific, CA). This was followed by c-DNA synthesis (using SuperScript^TM^ III Reverse Transcriptase (1808085, Thermo Fisher Scientific, CA) and RT-qPCR. For this last step, a commercial array of 84 primer pairs for the amplification of typical EMT-related genes was used (RT² Profiler PCR Array, PAHS-090Z RT^2^ Profiler^TM^ PCR Array Human Epithelial to Mesenchymal Transition, Qiagen, Germany) and data were normalized and processed using the delta-delta Ct method in the Qiagen online data analysis platform (accessible at: https://www.qiagen.com/ca/shop/genes-and-pathways/data-analysis-center-overview-page/). New cultures were generated for each biological replicate. From the genes provided as housekeeping genes on the qPCR array plates, Beta-2-microglobulin (BM2) was chosen for data normalization as it showed most consistent CT values across all samples.

### Statistics

Data were normalized as follows: values in the control group were normalized to the mean control value across biological replicates; values in sample groups were normalized to their respective technical means. This allowed us to capture the variability associated with the control samples. Values are shown as the mean ± standard error of the mean and results were obtained from a minimum number of independent experiments of 3. All data were analyzed in GraphPad (GraphPad Prism version 7.00 for Mac OS, GraphPad Software, La Jolla California USA, www.graphpad.com). For comparing two test groups, an F-test was used to determine equal variance, followed by an unpaired t-test whenever equal variance could be assumed, and an unpaired t-test with Welch’s correction, whenever variances were assumed unequal. For testing differences between 3 test groups or more, a one-way analysis of variance (ANOVA) was used, followed by the Tukey post hoc test in cases of equal variance. All tests were two-tailed, and asterisks represent the following p-value cut-offs: * <0.05, ** < 0.005, *** < 0.0005, **** < 0.0001.

## Results and Discussion

### Design of GLAnCE manufacturing and cell loading process

We set out to establish a user-friendly cell culture platform to generate cocultures that mimic the spatial structure of the tumour-stroma interface and to use this platform to study interface-specific tumour cell phenotypes. Interfaces were created between CAL33 Head and Neck squamous cell carcinoma (HNSCC) cells and CAFs sourced from resected human tongue carcinoma tissue. We designed our culture platform to enable coculture of two cell populations embedded in two, initially segregated collagen hydrogel compartments (Fig. 1A). These two-phase hydrogels were structurally supported and maintained in channel-shaped depressions within a polystyrene (PS) sheet.

**Figure 1.**
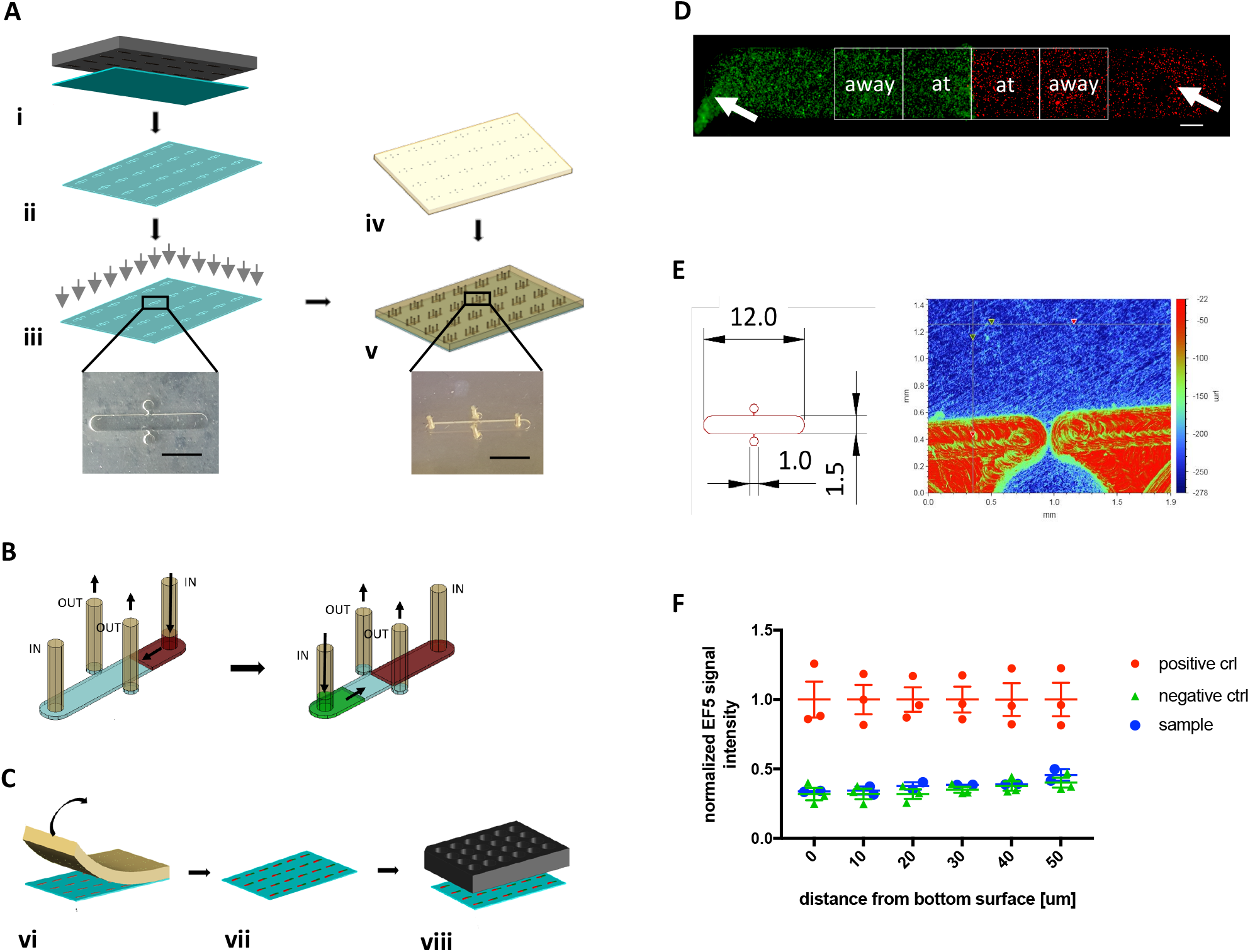
GLAnCE culture platform fabrication and generation of compartmentalized hydrogels. (A) Hot embossing (i) was used to imprint an array of open channels into a sheet of polystyrene (PS) (ii), which was subsequently UV-treated (grey arrows) to enhance its attachment towards hydrogels (iii); magnified image shows top-view of an individual ‘open channel’. Scale bar represents 5 mm. A thin silicone slab containing 1 mm through-holes, created using a bio punch (iv) was subsequently aligned with the PS sheet to create an array of temporarily ‘closed channels’ (v); magnified image shows top-view of an individual closed channel. Scale bar represents 5 mm. (B) Hydrogel compartment interfaces were generated within closed channels in a two-step injection process with a gelation step at 37°C between injections to enable gelation of each of the compartment phases; ‘IN’ labels injection ports, ‘OUT’ labels points for air escape. (C) Peeling off the PDMS slab returned channels from a closed to an open configuration (vi), thereby exposing the 6×4 array of compartmentalized hydrogels, (vii) which was then attached to a no-bottom, 24 well culture plate using an appropriately laser-cut, double-sided, polyacrylic tape (viii). (D) Fluorescent micrograph showing a full compartmentalized hydrogel immediately after seeding, containing tumour cell (red - mCherry) and CAF (green - eGFP) cell populations, which contact each other along a central compartment interface. ROIs of 2.25 mm^2^ (white boxes) show areas of interest for analysis, designated as ‘at’ and ‘away’ from the interface. Arrows indicate typical hydrogel injection artefacts (bubbles and areas of excess gel), which were systematically excluded from all analysis. Scale bar represents 500 μm. (E) Image of an individual, empty PS open channel obtained with profilometry showing a channel depth of approx. 250μm, with lower regions in the PS represented by colder colors. (F) EF5 intensity (a measure of local hypoxia) as a function of distance from the bottom surface of the plate through the thickness of the hydrogel layer. EF5 intensity in GLAnCE samples was not significantly different from negative controls (normoxia). The open channel-configuration of GLAnCE prevented the establishment of hypoxic gradients across the hydrogel after 5 days of culture, indicating that oxygen could freely permeate the thin tissue from the culture media, thereby removing the need for pumping systems. n=3.

We fabricated GLAnCE by hot embossing, using a custom-made aluminium stamp to impress a 6 x 4 array of channel-shaped depressions (termed ‘open channels’) into a sheet of virgin crystal PS (Fig.1A i, ii). PS channel depth remained consistent across the PS channel array after de-embossing from the aluminum stamp (i.e. no warping occurred), as confirmed by caliper measurements on two different stamp areas (randomly chosen at the sheet periphery versus centre) and on two channel areas (within versus outside a channel) (SI Fig. 1A). PS sheets were treated with UV light, in order to expose surface methyl groups (Fig. 1A iii). This was essential to enhance attachment of the collagen hydrogel to the PS well surface (48), which allowed to maintain the integrity of the hydrogels during subsequent steps in the fabrication process.

The UV-treated sheet of embossed PS was then sandwiched against a thin (~5mm) sheet of PDMS, which was modified, using a biopsy punch, to contain 2 inlet- and 2 outlet-holes per corresponding open channel on the PS sheet (Fig. 1A iv). Since clean PDMS is ‘sticky’ with respect to other smooth surfaces (largely because of Van-der-Waals bonds between both surfaces (52)), sandwiching resulted in reversibly ‘closed channel’ structures, which temporarily functioned as microfluidic channels with 2 inlets for cell loading and 2 outlets for air-escape (Fig. 1A v).

Next, cells were loaded into the closed channels in a step-wise procedure (Fig. 1B): First, fluorescently labelled carcinoma cells in a rat tail collagen type I suspension were injected into the right channel half through the right channel inlet (at a density of 5000 cells/channel). The volume of collagen solution injected determined the position of the hydrogel leading edge along the channel length. The collagen solution was gelled at 37 °C for 5 min. Next, a fluorescently labeled population of CAFs in collagen was injected from the left channel inlet (at a density of 10,000 cells/channel) until it reached the first, already gelled hydrogel compartment at the center of the closed channel. The air displaced by this process escaped through the two lateral outlet holes. This second collagen volume was gelled at 37 °C for 20 min. Rat tail collagen type 1 was chosen over bovine collagen type 1 (both are widely used in tissue engineering applications and *in vitro* tumour invasion models) for GLAnCE cultures, due to its characteristically faster gelation time (53), which prevented cell suspensions from settling at the bottom of the PS channels. This enabled a more homogeneous distribution of cells in the hydrogel in the z-dimension (SI Fig.1B). In order to confirm that the two-step hydrogel loading procedure yielded a truly continuous hydrogel slab with both sides in intimate contact, compartmentalized hydrogels were generated with gels containing secondary fluorescent antibodies (SI Fig. 1C). Upon gelation, each hydrogel side was fluorescently labelled in red or green and confocal imaging of the interface region showed that the fluorophores readily generated a gradient across the interface (yellow region), indicating that compartments were indeed in intimate contact with each other and no gap was present. The GLAnCE platform is also compatible with bovine collagen type I and other, commonly used hydrogels, such as Matrigel (SI Fig. 1D), as well as with different cell types (SI Fig 1E): CAL27 tongue carcinoma cells showed increased strand formation in CAF cocultures, similarly to CAL33 tumour cells (SI Fig 1Ei); coculture of MDCK cells, which are transformed, but not tumorigenic epithelial cells, with CAFs (SI Fig 1Eii) showed that CAFs could induce invasive behaviour in these initiated epithelial cells, an observation in line with early studies investigating pro-tumorigenic potential of fibroblasts on pre-neoplastic epithelia (54,55). Furthermore, we could also generate compartmentalized gels of dual stiffness (SI Fig. 1F).

After full gelation of both hydrogel compartments, the PDMS sheet was peeled from the PS sheet, exposing the hydrogels, which remained immobilized inside the open-channels (Fig. 1C vi, vii). Finally, we used a custom-cut, double-sided polyacrylic tape to attach the PS sheet containing the gel-filled channels to a bottom-less 24 well plate. Note that we confirmed that the presence of the tape did not impact cell viability (SI Fig. 1G). Overall, this workflow produced a 24 well culture plate with one compartmentalized hydrogel at the bottom of each individual well (Fig. 1C viii).

Cells in GLAnCE culture grew uniformly across the hydrogel’s length, as assessed using an MTT assay after 5 days of culture and were metabolically active everywhere, highlighting the uniformity of the hydrogel environment generated using our seeding procedure (SI Fig. 1H). Cell numbers for GLAnCE cocultures were carefully optimized: for tumour cells, a high density was chosen, which would still allow for visual identification of individual aggregates and any changes to aggregate morphology after 5 days of culture. For CAFs, the specific seeding density used directly reflects the critical amount of fibroblasts necessary for eliciting phenotypic changes in tumour cells at the interface in a reproducible fashion.

The geometry of the channels was selected such that the channel length was sufficient to include distinct hydrogel zones (‘at’ the interface, ‘away’ from the interface), and so that regions immediately underneath the hydrogel loading inlets could be systematically excluded from all analyses, as these presented injection artefacts such as decreased cell density (bubbles) or increased cell density (excess collagen-cell suspension) (Fig. 1D and Fig. 1E, left). Furthermore, the compartment interface was perpendicular to the long channel axis. This allowed quantification of tumour-stroma interface movement over longer distances, while still minimizing the amounts of cells and collagen volume required. We restricted channel depth to 276μm +/-31 µm (Fig. 1E, right), to ensure cells would experience an environment thick enough to remain in 3D, while also being thin enough for easy imaging of live samples. Because GLAnCE hydrogels were maintained in open PS channels, culture medium could access the entire gel surface and cells therefore did not experience significant gradients of oxygen (unlike the gradients often present in thicker 3D culture setups). We assessed the presence of hypoxia throughout the hydrogels using EF5 staining (Fig. 1F) and detected no hypoxic gradients in the hydrogels’ z dimension, nor within individual tumour cell aggregates.

In the resulting microtissue, CAFs and carcinoma cells were completely segregated, but contacted each other along the central compartment interface (Fig. 1), reminiscent of tissue architecture prior to tumour-stromal mixing at the invasive tumour front. Over five days in culture we observed remodeling of the interface region and mixing of epithelial and stroma cell populations (Fig. 2A). Due to the manual loading process used to generate GLAnCE hydrogels, the shape of the compartment interfaces was found to be either straight or curved, in approximately 45% and 49% of interfaces examined, respectively. The remaining 6% of interfaces were irregular in shape. Neither of these interface shapes had a significant impact on the extent of interface movement over time (SI Fig.1I). The GLAnCE design therefore provided a robust platform to establish compartmentalized cocultures and to study the movement of the tumour-stroma interface.

**Figure 2.**
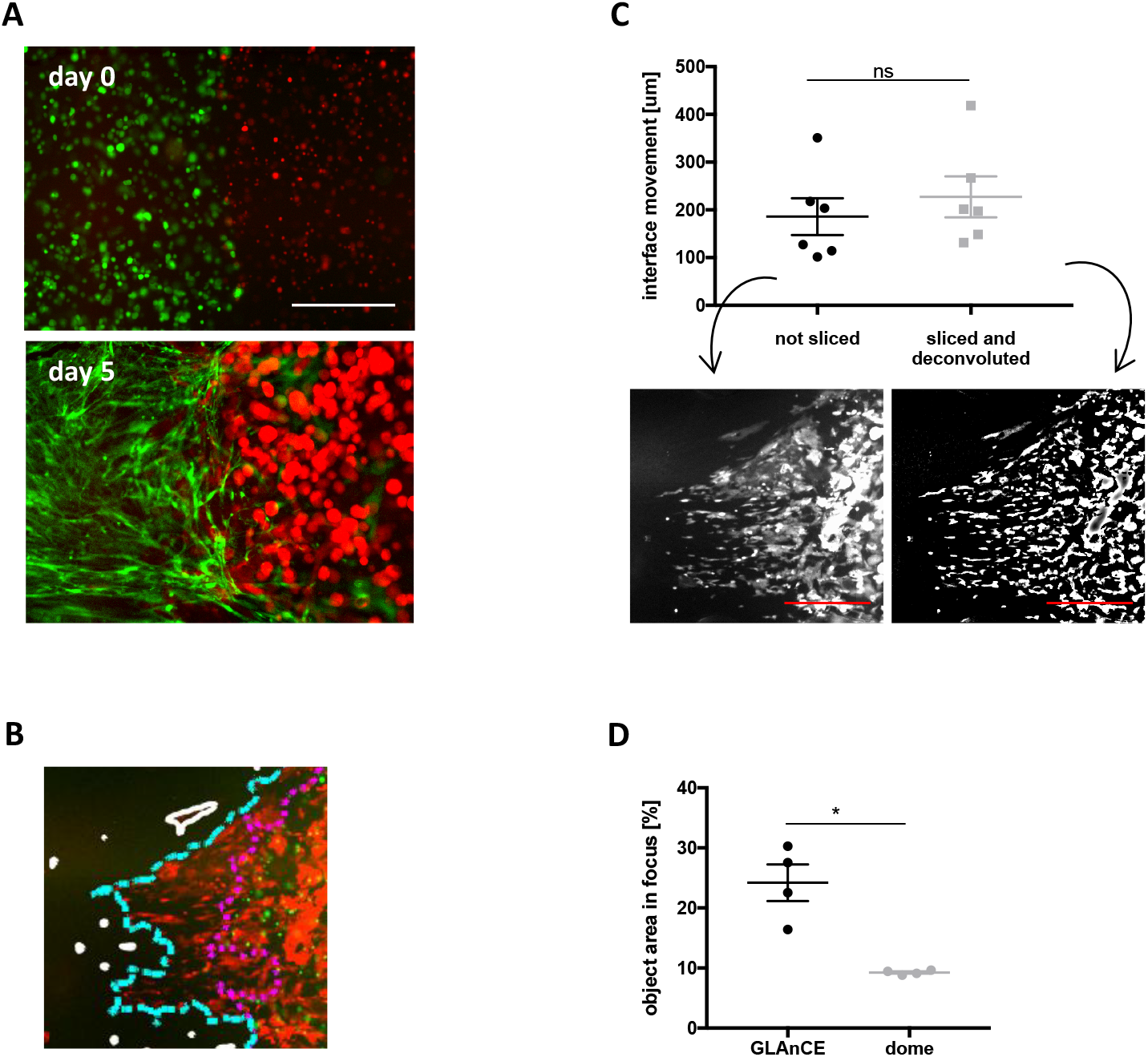
GLAnCE is compatible with widefield microscopy and enables improved object visualization. (A) Widefield fluorescence images of interface remodeling in GLAnCE from day zero (day of hydrogel seeding) until day 5 in coculture between tumour cells (red – mCherry) and CAFs (green - eGFP). Scale bar is 500 μm. (B) Overlay of tumour cells at day 0 (green) and at day 5 (red). The automated algorithm detects the tumour compartment interface at day 0 (purple) and at day 5 (blue). 2D tumour cell clusters distant from the interface (which are omitted from the analysis) are outlined in white. The distance is then quantified between the purple and blue lines at each y coordinate to calculate a mean interface displacement. (C) Widefield fluorescent microscopy was as effective at detecting individual cells moving outwards from the compartment interface, as multi-slice acquisition followed by image deconvolution. The mean interface movement from 6 GLAnCE samples was quantified using our automated algorithm for images obtained using both imaging approaches. Example images of the same sample acquired using each imaging strategy are shown. Tumour cell fluorescent signal is in white, scale bar is 500 µm. Error bars represent SEM, n=6. (D) The proportion of object area in focus in micrographs of GLAnCE hydrogels vs. hydrogel domes (another high-usability hydrogel culture setup) was assessed using fluorescent beads of uniform size. Hydrogel geometry in GLAnCE resulted in a significant decrease of blurry objects (object areas out of focus), due to the restricted z-dimensionality of the hydrogels. Error bars represent SEM, n = 4.

### Quantification of tumour cell invasion using widefield microscopy

Having optimized our device fabrication process, we next wanted to demonstrate key design advantages of our platform for improved usability. Firstly, we considered the mode of data collection, which needed to be compatible with live imaging of CAF and tumour cell movement away from the initial interface location (Fig. 2A and SI Fig. 2A). We quantified carcinoma cell position with respect to the interface (as a measure of tumour cell movement from the interface) using a custom-built automated MATLAB (Mathworks) algorithm. The algorithm automatically detects the outline of the compartment interface at day 0 and at day 5, and then calculates the mean change in compartment interface position and the area associated with this change (Fig. 2B). Of note, this approach was tailored to tumour cells moving collectively, with preserved cell-cell adhesions; in fact, our algorithm was not able to accurately localize compartment interface outlines of tumour cells moving within GLAnCE as single cells, which therefore represents a limitation of this image-based strategy for the quantification of cell movement. We assessed the performance accuracy of our automated algorithm by benchmarking automated measurements against manual interface tracing carried out by 3 independent blinded users. The automated interface detection algorithm produced results within the variability range of manual measurements (SI Fig. 2B).

Visualization of cell movement within the 3D hydrogel space was possible using both confocal and widefield fluorescence imaging. We reasoned that the use of wide-field microscopy would be advantageous in terms of reducing the time required for tissue imaging. To determine if wide-field fluorescence imaging was in fact suitable for data acquisition, we compared the fluorescence signal detected (as measure of cell location) in single-slice versus multi-slice images (optically sliced in the z-dimension) (Fig. 2C). The latter set of images were deconvoluted to remove out-of-focus blur. Cells that moved the furthest away from the bulk compartment had the lower fluorescence, compared to the bulk. Visualization of these cells was thus used as a surrogate measure of fluorescence detection capacity in both imaging modalities. Similar spatial patterns of fluorescence signal were detected using single-slice and multi-slice images and no statistical difference was observed between non-sliced and sliced + deconvoluted images at 4X magnification (the magnification required to visualize interface movement in our system). This suggested that optical image slicing was not necessary, in order to visualize even small clusters/individual cells within the compartmentalized hydrogels.

We also assessed the ability of GLAnCE to maximize the number of objects in focus within a given focal plane. When we compared the GLAnCE hydrogel geometry to hydrogel dome cultures (a standard culture method for cell aggregate and organoid culture) seeded at the same cell densities, we found a greater area in micrograph occupied by objects in focus in GLAnCE (Fig. 2D), indicating that more aggregates were located at the same z-position, thereby aiding visualization of aggregate morphologies across the hydrogel.

### Movement of the compartment interface results from CAF-induced carcinoma cell proliferation

Having validated our analysis methods in GLAnCE, we next set out to quantify the impact of CAF coculture on the remodeling of the tumour-stroma interface. To do this we assessed interface movement in mono- and cocultures and observed that tumour cells moved significantly more in coculture compared to monoculture (Fig. 3A and Fig 3B i). Furthermore, tumour cells appeared to move homogenously outwards from the interface (Fig 3A). The area occupied by the moving tumour cells was also larger in coculture compared to in monocultures. Note that during hydrogel gelation in GLAnCE, a small fraction of tumour cells inevitably contacted the bottom channel surface. These cells moved as small, 2D cell clusters and were easily identifiable in micrographs. These cells were excluded from quantification as their inclusion concealed any differences between mono- and cocultures, likely due to migration of these cells being influenced through contact with the stiff PS surface (SI Fig.3A). This co-culture induced tumour cell invasion was observed in compartmentalized cocultures with 3 distinct, patient-derived CAF lines (SI Fig. 3B). Note that all 3 CAF lines resulted in morphological changes in tumour cell aggregates at the compartment interface, indicative of HNSCC CAF pro-invasive activity, however, only CAFs sourced from patient 1 (CAF line number 61137) were able to induce a consistent and significant tumour interface movement. This observation highlights the heterogeneity of the pro-proliferative capacity between CAF populations. Because of their combined effect on tumor cell proliferation and invasion, CAFs from patient 1 were utilized for all further experiments.

**Figure 3.**
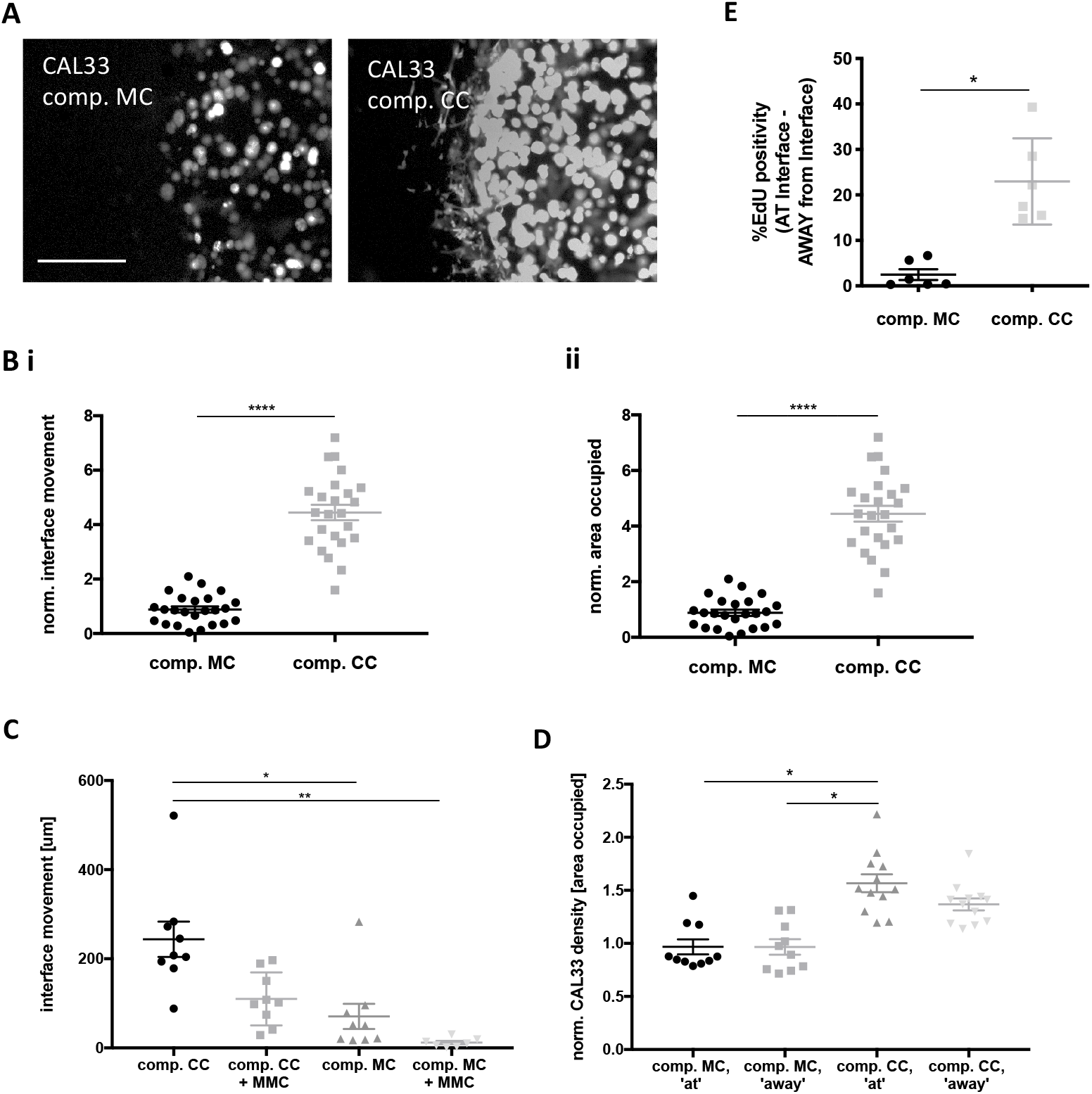
Tumor cell movement from the interface is enhanced by CAF-secreted signaling factors. (A) Representative images of tumor compartment interfaces (white) after 5 days in compartmentalized monoculture (comp. MC) and compartmentalized coculture (comp. CC). Scale bar represents 500 μm. (B) Interface movement over 5 days in culture was significantly higher in CAF cocultures. Similarly, normalized area occupied by the moving tumor cells was also higher in CAF cocultures. Error bars represent SEM, n= 10. (C) MMC-treated compartmentalized cocultures showed significantly decreased interface movement, compared to control cocultures, revealing the contribution of proliferation to compartment expansion. Error bars represent SEM, n=3. (D) Area coverage of tumor cell aggregate fluorescence was quantified at and away from the interface as a measure of tumor cell proliferation. Compared to monoculture, tumor cell aggregates at the compartment interface in coculture occupied the largest area, suggesting very localized CAF-secreted growth factors or direct interaction with CAFs as the pro-proliferative signals. Data were normalized to day 0 cell density. Error bars represent SEM, n=3. (E) Tumor cell proliferation was assessed by EdU staining at and away from the compartment interface in mono and co-cultures and the difference between regions was higher in CAF comp. co-cultures. Error bars represent SEM, n=2.

We next wanted to demonstrate that we could use GLAnCE to probe the mechanisms behind the outwards movement of the interface, and specifically the relative contributions of tumour cell proliferation versus active invasion. When we pre-treated tumour cells with 10 μM mitomycin C to block cell proliferation, we observed significantly decreased interface movement in both co- and monocultures. While inhibition of proliferation did not fully abolish interface movement, this data suggest that tumour cell proliferation is a major contributor (Fig. 3C). This approach further showcases the use of GLAnCE to assess the effect of phenotype-altering compounds on both cell populations in their spatial interdependence.

To identify whether the pro-proliferative signal from the CAFs was localized at the interface or whether it impacted the entire tumour compartment, we quantified the percentage area covered by tumour cell fluorescence (a metric of cell growth) within regions of interest both at and away from the compartment interface in mono- and cocultures (Fig. 3D). We observed a greater, but not statistically different, carcinoma cell area ‘at’, compared to ‘away from’ the interface in CAF cocultures. Aggregate size at the interface in cocultures were also significantly more populated with tumour cells than either location in monoculture, suggesting that the CAF pro-proliferative effect was mediated by a direct interaction with CAFs (Fig. 3D). This was in agreement with EdU staining of tumour cell monocultures and compartmentalized cocultures (Fig. 3E) where tumour cell division was highest at the interface in compartmentalized cocultures, highlighting the pro-proliferative role of direct tumour cell-CAF interactions. In fact, treatment of tumour cell monocultures with conditioned media from both CAFs and CAF-tumour cell cocultures failed to significantly increase interface movement (SI Fig. 3B). Note however that conditioned media was prepared with media from CAFs and CAF-tumour cell cultures and that this approach may therefore not have captured matrix-bound signaling molecules, nor soluble factors smaller than 3kDa (the size corresponding to media concentration columns filters used).

### CAFs mediate tumour cell dissemination from the interface through invasive strand formation and single cell dispersal

In addition to increased interface movement via proliferation, the GLAnCE platform allowed us to visualize and quantify morphological changes to the tumour cell aggregates indicative of an invasive phenotype. Tumour monocultures contained proliferating tumour cell clusters with near-circular perimeters, irrespective of their location within the gel (Fig. 4A, top panel). By contrast, tumour cells in compartmentalized coculture with CAFs formed organized strand structures protruding from aggregates, specifically at, but not away from, the compartment interface (Fig. 4A, middle panel). When we quantified strand morphology of single-aggregates in compartmentalized cocultures, we found that tumour cell clusters at the interface deviated significantly from a circular perimeter, while clusters away from the interface displayed circle-like morphologies identical to those observed in monocultures (Fig. 4B).

**Figure 4.**
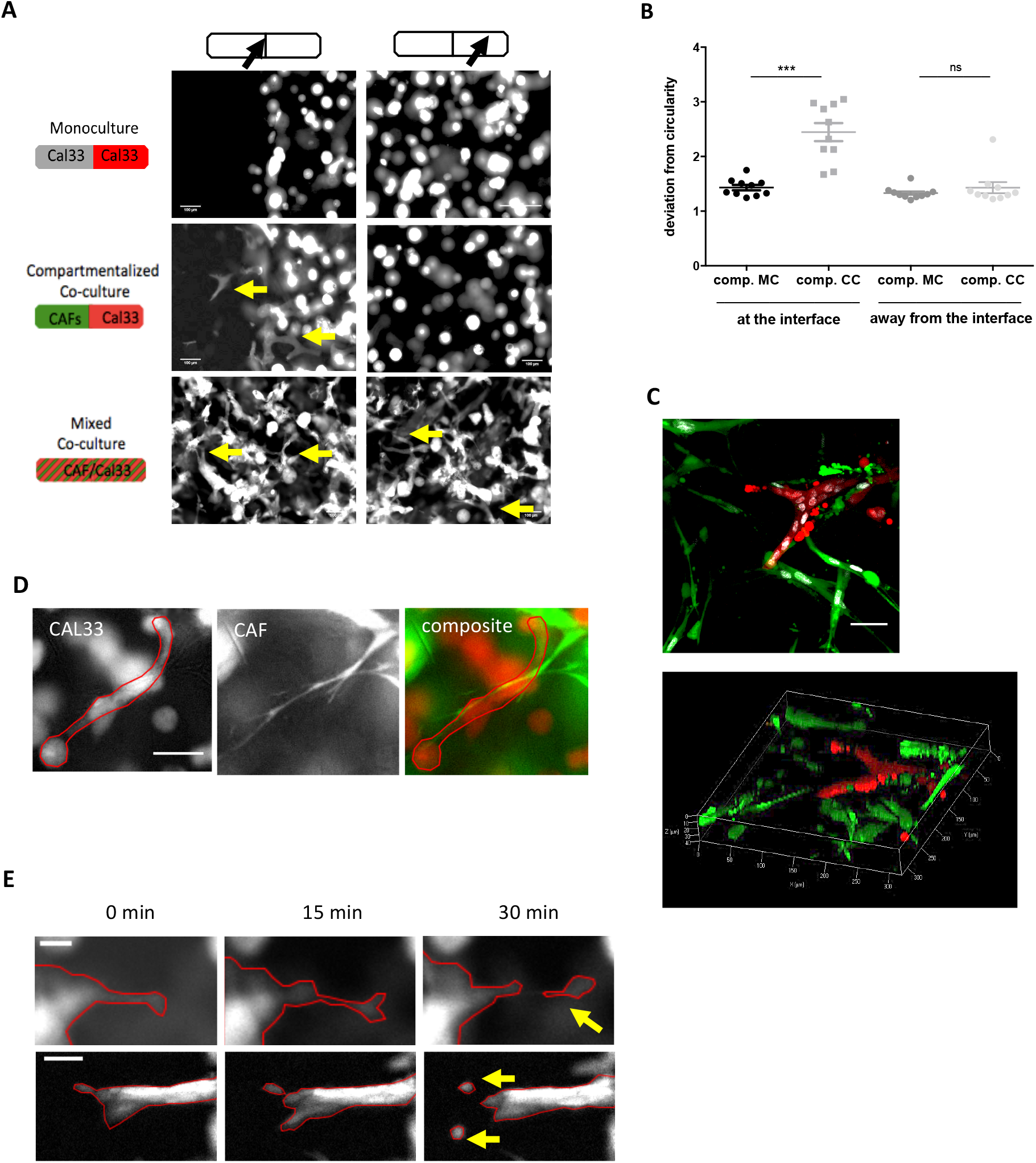
Tumour strand formation is CAF-dependent and leads to carcinoma dissemination as single cells. (A) Tumour cell invasive phenotype was dependent on the presence of CAFs at the compartment interface. Arrowheads point towards tumour stand structures, scale bar represents 100 μm. (B) Tumour cell morphology was consistent with invasive aggregate elongation at, but not away from, the compartment interface. Morphology was quantified as deviation from circularity, which approaches 1 for near-circular objects and is increasingly higher, the more elongated the object. Error bars represent SEM, n = 3. (C) Confocal stack and 3D rendering of tumour invasive strand shown in (A), confirming tumour cell protrusions (red) were indeed structures protruding into the three-dimensional hydrogel-CAF (green) space. Scale bar represents 50 μm. (D) CAFs and tumour strands (traced in red for clarity) aligned and were in immediate contact with each other, with CAFs oriented along the long strand axis. Scale bar represents 100 μm. (E) Representative images of tumour strands (traced) giving rise to independently invading, single cells from their tips, with arrows pointing towards mesenchymal (top) and ameboid (bottom) invasive morphologies. Scale bars represents 20 μm (top) and 50 μm (bottom).

The observed tumour cell strand structures were multi-cellular and extended in three-dimensions (Fig. 4C). Furthermore, strands were consistently accompanied by CAFs oriented in parallel and located immediately adjacent to the direction of strand elongation (data not quantified, Fig. 4D) suggesting CAFs were required for strand formation. Indeed, strand formation from tumour aggregates occurred throughout the entire carcinoma hydrogel compartment when tumour cells and CAFs were homogeneously mixed (Fig. 4A, bottom panel). In addition, tumour cell aggregates in monoculture did not form strands in conditioned media from CAF monocultures or CAF-tumour cell mixed cocultures (SI Fig. 4), further suggesting that CAFs were required to facilitate strand formation.

In monoculture, tumour cells grow in dense multi-cellular clusters, which increase in size over time, resulting in a sharp decrease in the ECM space available between aggregates. In contrast, CAFs are present as single cells and created an environment with plenty of space to accommodate additional tumour cells as they moved outwards from the compartment interface. We therefore wanted to assess whether strand assembly in tumour monocultures could be prevented simply by a lack of free space. To do this we generated hydrogels in which the tumour compartment was interfaced with an acellular collagen compartment (which, similar to CAFs, provided unrestricted space for tumour aggregate growth at the interface). Aggregate morphology in these gels with one acellular gel compartment were indistinguishable from those containing tumour cells in both compartments (data not shown). Together these data strongly suggest that direct contact with CAFs or CAF-mediated remodelled matrix, as opposed to soluble factor paracrine signalling or lack of free space, were responsible for tumour strand formation in compartmentalized coculture.

Tumour cells are known to use different invasion mechanisms, as indicated by cell morphology, depending on the extent of tumour cell local confinement, whereby greater confinement promotes collective migration (56). Time-lapse imaging of GLAnCE allowed us to further explore the invasive tumour cell morphologies in strand structures at the interface. Specifically, we observed both collective carcinoma cell migration to form the strands and individual carcinoma cell invasion at the tips of already formed strand structures (Fig. 4E, SI Movies 1 and 2). Cells exhibiting both elongated mesenchymal morphology (as seen in Fig. 4E, top panel) and spherical amoeboid morphologies (Fig. 4E, bottom panel) were seen to emerge from strand tips. We speculate that we observed this spectrum of invasive behaviours as a result of mounting remodeling and breakdown of local matrix confinement at the interface, due to increasing levels of infiltrating CAFs over time, which established a matrix conducive of, first, collective tumour cell invasion in strands, and subsequently invasion as individual cells with mesenchymal, and even amoeboid, morphologies as more matrix space becomes available.

### Probing the impact of matrix properties on CAF-enhanced tumour cell invasion

We next set out to use GLAnCE to explore in more detail the mechanism by which CAFs induced tumour cell strand formation and invasion at the interface. CAF-induced invasion could result from remodeling of the matrix to create migration permissive tracks for tumour cells, and/or from induction of epithelial-to-mesenchymal transitions (EMT) (57). We first assessed the impact of the extracellular matrix properties on CAF-induced invasion. The properties of the tumour ECM have been shown to be pivotal in regulating tumour aggressiveness *in vivo* and typically, more porous matrix structures, are associated with an increase in invasion (56).

We attempted to assess tumour invasion into CAF-remodelled, and subsequently decellularized matrices, as others have done (58); however, GLAnCE hydrogels, due to their thinness, were too fragile for this approach. Therefore, to test whether loss of matrix confinement, rather than direct cell-cell interaction with CAFs, was sufficient to allow strand formation and invasion in our system, we quantified strand formation in collagen matrices of different mesh-sizes. Previous work has shown that bovine collagen type 1 is characterized by a non-stabilized mesh structure of collagen fibrils, resulting from the lack of telopeptides, which are removed in the process of protein extraction from the primary bovine tissue. Tumour cells have been shown to move within this matrix without the need for MMP-dependent matrix proteolysis (53,59). In GLAnCE, strand structures were detected at the CAF-tumour interface in all hydrogel matrices containing stabilized/native (i.e. rat tail) collagen type I, provided CAFs were present (Fig. 5A, top). By contrast, in non-stabilized (i.e. bovine) collagen type I, strands were observed throughout the tumour compartment (not just at the interface) in mono- and cocultures (Fig. 5A, bottom and Fig. 5B). Further, where strands were observed, morphological analysis revealed the expected deviations from circularity (a proxy for strand elongation) (Fig. 5C). We speculate that the large mesh size characteristic of the bovine collagen enabled CAF-independent tumour invasion. This data suggests a mechanism by which CAFs initiate invasion of tumour cells by generating spaces within the stabilized/native collagen matrices at the tumour-stroma interface in our platform.

**Figure 5.**
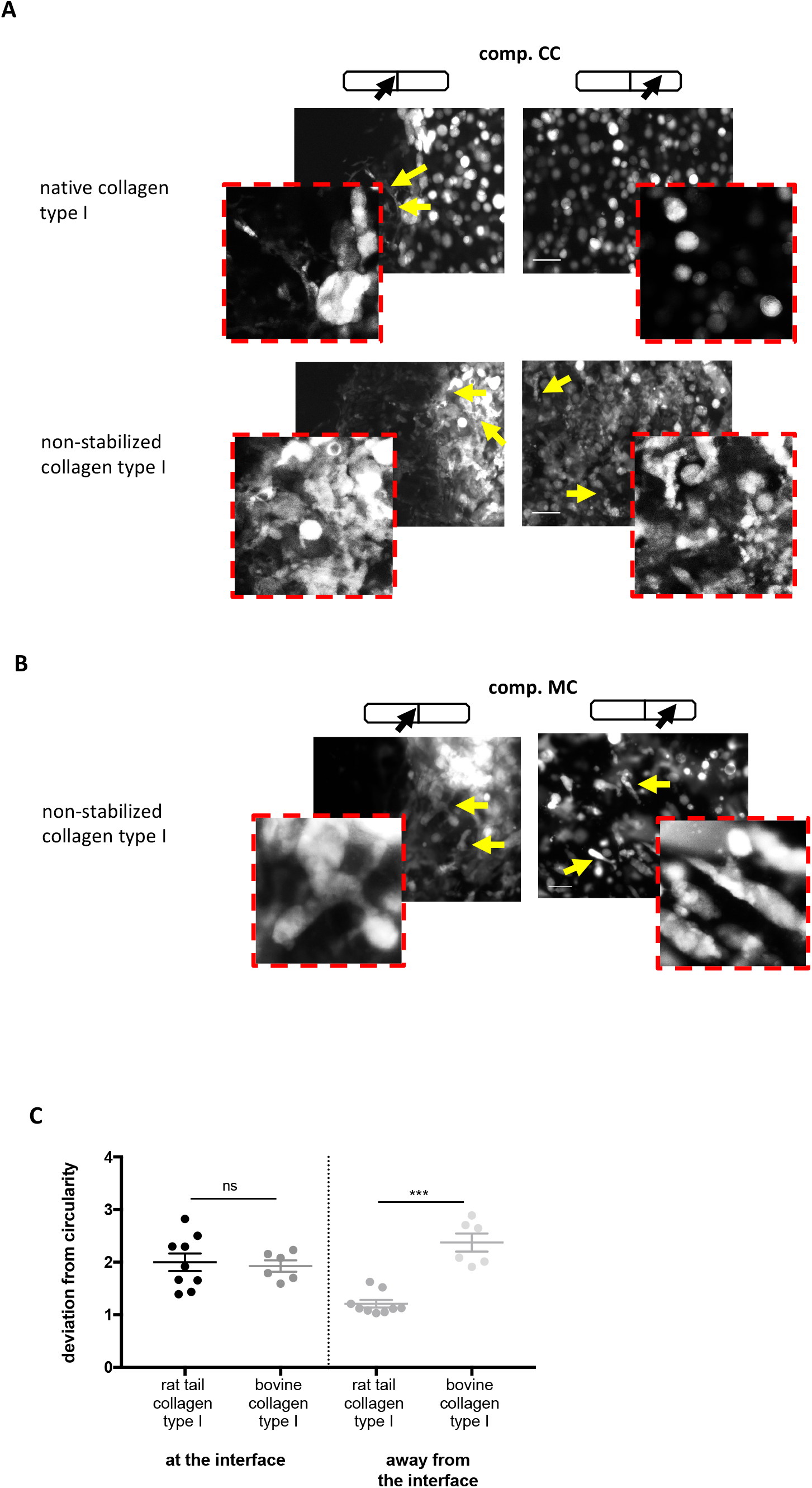
Hydrogel composition and stiffness affects tumour compartment expansion and invasiveness. (A) Tumour cells cultured in compartmentalized cocultures (comp. CC) with CAFs in matrices of different compositions. Tumour cells gave rise to strand structures exclusively at the interface in native (rat tail) collagen (top panel), while in non-stabilized (bovine) collagen matrices (bottom panel) they produced strands throughout the gel. (B) Tumour cells also displayed invasive capacity in bovine compartmentalized monoculture (comp MC), ubiquitously throughout the hydrogel. Insets in A and B show individual strand structures. (C) Invasive morphology of tumour strands in different matrices was quantified at and away from the compartment interface. Error bars represent SEM, n= 3.

### Analysis of compartment interface-specific versus long range CAF-tumour cell interactions shows KRT14 upregulation at the invasion-permissive compartment interface

Movement of tumour cells into free spaces within an invasion-permissive matrix requires changes in tumour cell motility and acquisition of mesenchymal properties (57). This epithelial-to-mesenchymal transition (EMT) occurs in response to both paracrine signals secreted by CAFs, as well as direct contact with CAFs or CAF-remodelled matrix (24,60). To identify the genes up-regulated in tumour cells at the tumour-stroma interface involved in EMT, we performed an RT-qPCR analysis using a set of 84 primer pairs for the amplification of typical EMT-related transcripts (RT² Profiler PCR Array). To differentiate transcriptional changes in tumour cells in response to signals at the invasion-permissive matrix tumour edge versus those induced in the bulk tumour compartment by long-range signaling originating from CAFs, we created compartmentalized gels in three distinct culture arrangements: compartmentalized tumour cell monoculture (comp. MC), tumour cell-CAF compartmentalized coculture (comp. CC), and mixed coculture (mixed CC), where both cell populations were homogeneously mixed before seeding into the device (the total cell numbers were kept constant) (Fig. 6A). Carcinoma cells in fully mixed cultures represented the interface zone where stromal content was high. Carcinoma cells in compartmentalized coculture contained mainly tumour cells in the bulk with only a small fraction of tumour cells located at the interface in direct contact with the CAFs or CAF remodelled matrix. We reasoned that this sample would allow us to determine the effect of long-range signaling from the CAFs on the bulk tumour compartment. A comparison of these two groups to monoculture would enable us to isolate the transcriptional changes specifically associated with the interface zone.

**Figure 6.**
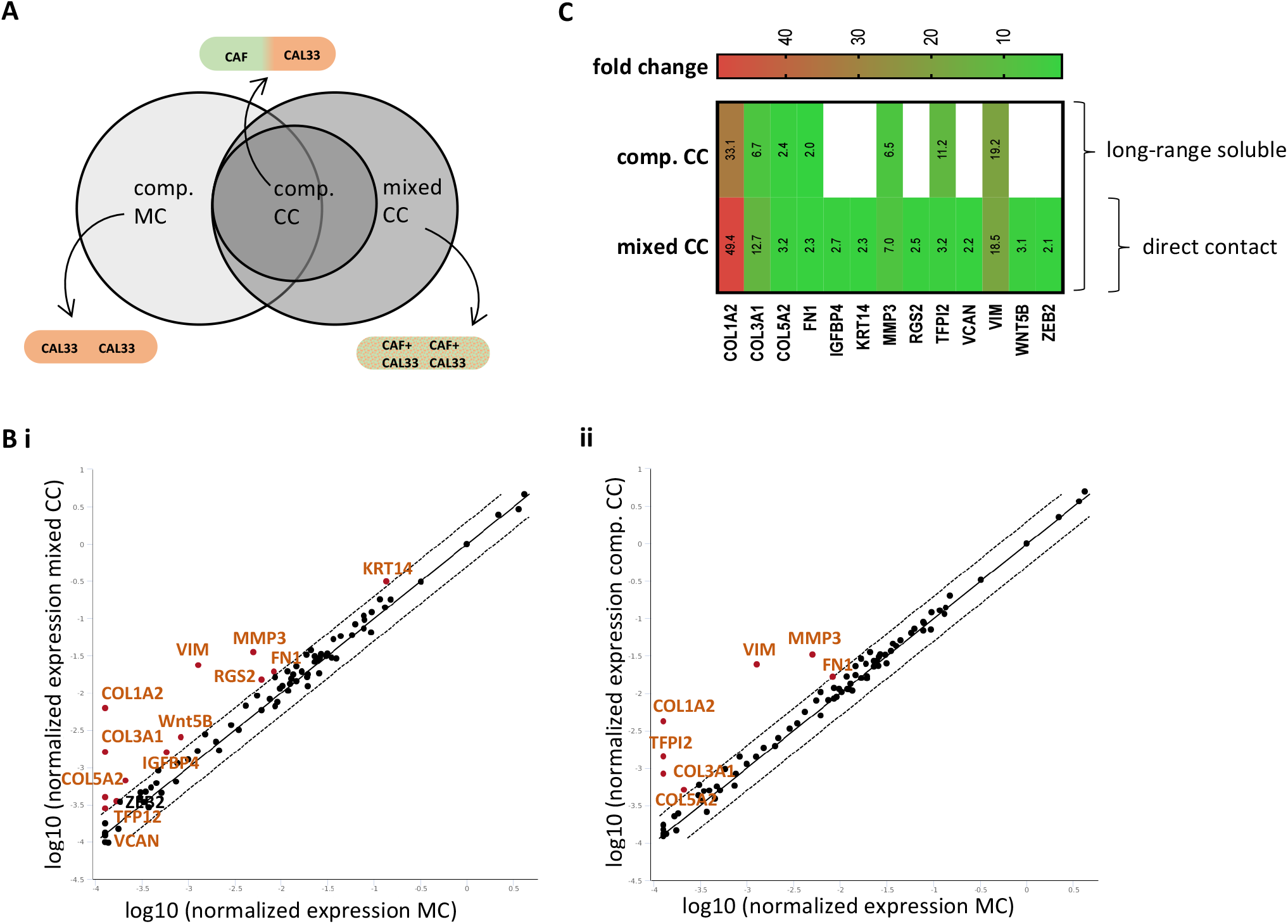
Expression of KRT14 is induced at the *in vitro* tumour front. (A) Experimental design showing how compartmentalization in GLAnCE can be utilized to discern between tumour-stromal interface-specific, pro-invasive signals and soluble signaling that affects the bulk of the tumour compartment. MC and CC are monoculture and coculture. (B) Gene expression correlation plots showed an upregulation of several EMT-related genes (orange) in mixed cocultures (mixed CC) (i) and compartmentalized CC (comp. CC) (ii), compared to ctrl (comp. MC). Central range indicates a fold regulation cut-off of +/- 2, i.e. unchanged gene expression. (C) The differences between CC groups were a direct result of the amount of invasion-permissive or CAF-rich matrix within the hydrogel and was used to identify genes differentially expressed in an invasion-permissive environment with CAFs. Warmer colours indicated higher fold-regulation changes.

Cells were harvested from GLAnCE cultures by enzymatic matrix digestion, tumour cells were then isolated via fluorescence-activated cell sorting (FACS) (Si Fig. 5) and EMT gene transcription was assessed using a RT² Profiler PCR Array. As expected, significant transcriptional upregulation of EMT genes was observed in both coculture configurations, compared to tumour cell monoculture. Further, genes from compartmentalized cocultures (characterized by minimal CAF content at the interface) represented a subset of all genes upregulated in mixed coculture (characterized by much higher CAF content) (Fig. 6A and Fig. 6B). These transcriptional changes present in both coculture groups likely represented responses to secreted signals from CAFs, that affected tumour cells across the entire tumour compartment (Fig. 6C). For example, we observed an upregulation of matrix-metalloproteinase 3 (MMP-3) in both coculture configurations. MMP-3 is known to cleave collagen type IV (61) and laminin (62), and to initiate a positive MMP-3 self-activation loop (63); upregulation of MMP-3 in tumour cells throughout the hydrogel in response to secreted CAF signalling from the stromal compartment is consistent with our observations that tumour cells formed invasive structures across the entire tumour compartment when seeded in coculture in Matrigel (which is rich in laminin) (SI Fig. 1C). Interestingly, while these EMT-related changes were elicited by long-range CAF secretions, matrix-confinement in collagen still prevented tumour cell invasiveness in CAF-devoid matrix regions. Thus, this cohort of EMT markers could potentially represent transcriptional changes that preceded those driving the invasive phenotype at the compartment interface. This observation highlights that while changes in tumour cell gene expression occurred in response to CAF secretions, on a functional level these gene changes did not correlate with, and were therefore likely by themselves not sufficient for inducing, the invasive phenotype.

A separate cohort of genes including Wnt 5B, ZEB2, Versican (VCAN) and cytokeratin 14 (KRT14), was upregulated in mixed cocultures exclusively, where all tumour cells experienced the invasion-permissive environment (with high CAF content) throughout the hydrogel (Fig. 6C). This was in contrast to compartmentalized hydrogel samples, in which only a small fraction of tumor cells experienced the signals of an invasion-permissive environment and thus did not show significant upregulation of this gene cohort. Aberrant Wnt signalling is characteristic of many types of squamous cell carcinoma (64). In a study investigating the role of Wnt signalling on the metastatic potential of oral SCC, WNT5B signalling was required for carcinoma cells migration through active CdC42 and RhoA signalling (65). ZEB2 is a repressor of E-cadherin expression (66), is upregulated in tumour cells, and leads to EMT-like changes in tumour cell phenotype (67). VCAN, an ECM proteoglycan known to promote most aspects of tumour progression, including proliferation, differentiation, migration and adhesion, in several cancers (68), has been reported to localize preferentially in the vicinity of the stromal compartment and specifically in proximity to vasculature (69).

Most notably, tumour cells present within an invasion-permissive matrix with high CAF content upregulated their expression of KRT14 (Fig. 6C). KRT14 is a marker of the proliferating basal progenitor cell population in epithelial tissues that are in contact with the underlying ECM (70). KRT14 expression is upregulated at the tip of invasive strands in breast cancer 3D *ex-vivo* assays and *in vivo*, where KRT14^+^ ‘leader cells’ facilitate collective tumour cell invasion and metastases (71,72). In HNSCC specifically, KRT14 expression has been documented as a marker of micro-metastases (73), and is abundant in primary tumours (74). Of note is the co-expression of vimentin and KRT14 in our system, a pattern classically not defined as EMT, which however has been observed in early oral SCC samples and points towards a retention of the tumour cells’ epithelial phenotype, even while acquiring mesenchymal traits, as reported by others (75). In order to assess the generalizability of CAF-induced overexpression of this gene in invasion-permissive environments KRT14 expression was further quantified in CAL27 tumour cells, another HNSCC cell line (SI Fig. 5B). As was observed in CAL33 carcinoma cells, these tumour cells upregulated their expression of KRT14 in mixed coculture with CAFs (mean fold change > 2), suggesting that cytokeratin 14 may be a mediator or CAF-induced EMT in HNSCC cell lines, an observation that warrants further investigation.

Overall, EMT was potentially engaged by tumour cells in GLAnCE upon establishment of an invasion-permissive matrix by CAFs, though a direct distinction between the effects of a CAF-remodelled matrix and direct contact between CAFs and tumour cells will require further investigation.

## Conclusions

Numerous 3D *in vitro* culture platforms are available for exploring the impact of coculture and cell-matrix interactions on tumour cell phenotype. However, only a small subset of these platforms provide the control to impart a tissue-reflective architecture and to spatially prime the interactions between two cell populations (46,76). Furthermore, many of the models within this subset are challenging to use, resulting in low adoption of complex architectural tumour-stroma models by the wider cancer biology community. In an attempt to address this issue, we have created GLAnCE, a culture platform characterized by i) a device assembly method based on transiently closed channels that is therefore exempt from typical channel-bonding procedures, ii) hydrogel structures open to culture media, which removes the need for pumps and perfusion to counteract the generation of oxygen gradients, and allows straight-forward harvesting of cells from the channels, and iii) image-based data acquisition that does not rely on optical slicing and is non-sacrificial, thus saving significant time and enabling time-lapse experiments of the entire hydrogel array without incurring significant differences in time between the first and last image acquired across the hydrogel array. The GLAnCE culture platform allows users to study stromal and epithelial tumour interaction dynamics specifically by enabling visualization and quantification of the spatial-temporal dynamics of disease progression during invasion, without being limited to bulk measurements in traditional mono or cocultures.

Using GLAnCE, we have demonstrated that tumour compartment interface movement relies on tumour cell proliferation and that CAFs exert localized pro-proliferative effects on carcinoma cells. We also observed CAF-mediated pro-invasive effects specifically at the tumour-stroma interface, which resulted in carcinoma cell strand formation and tumour cell dispersal from the tips of these invasive strands. CAF-induced tumour cell invasion was likely due to both remodeling of the extracellular matrix to render it invasion permissive and concurrent changes in EMT-related gene transcription in the tumour cells. For example, tumour cells upregulated their expression of KRT14 at the tumour-stroma compartment interface. Notably, while heightened KRT14 expression at the epithelial-stromal border has been observed *in vivo* (71,77), this study explicitly implicates CAFs as inducers of this expression, either directly or via establishing an invasion-permissive matrix environment.

The GLAnCE platform enables quantification of CAF induced tumour cell phenotypes with spatial and temporal resolution. Furthermore, the design of the platform has been optimized for easy adoption by the broader research community in terms of usability. Importantly, the GLAnCE platform is compatible with various tumour cell types and fibroblasts. GLAnCE therefore offers the potential for probing spatial and temporal interactions between two cell populations and for modeling a range of diseases in regenerative biology.

## Acknowledgements

This work was funded by a Natural Science and Engineering Research Council (NSERC) and Canadian Institute of Health (CIHR) Collaborative Health Research Program (CHRP) grant (FRN 146469) to APM, an Ontario Trillium Scholarship and a NSERC CREATE M3 fellowship to ED, a NSERC CREATE M3 fellowship to JLC and a NSERC CREATE M3 fellowship to NCW.

## Data Availability Statement

The data that support the findings of this study are available from the corresponding author upon reasonable request.

**SI Movie 1:** Invasive strands (traced in green) giving rise to tumour cells moving independently from the strand tips. Scale bars represents 20 µm.

**SI Movie 2:** Invasive strands (traced in red) giving rise to tumour cells moving independently from the strand tips. Scale bars represents 50 µm.

