## Supplementary material for "Gels for Live Analysis of Compartmentalized Environments (GLAnCE): A Tissue Model to Probe Tumour Phenotypes at Tumour-Stroma Interfaces": Si figures

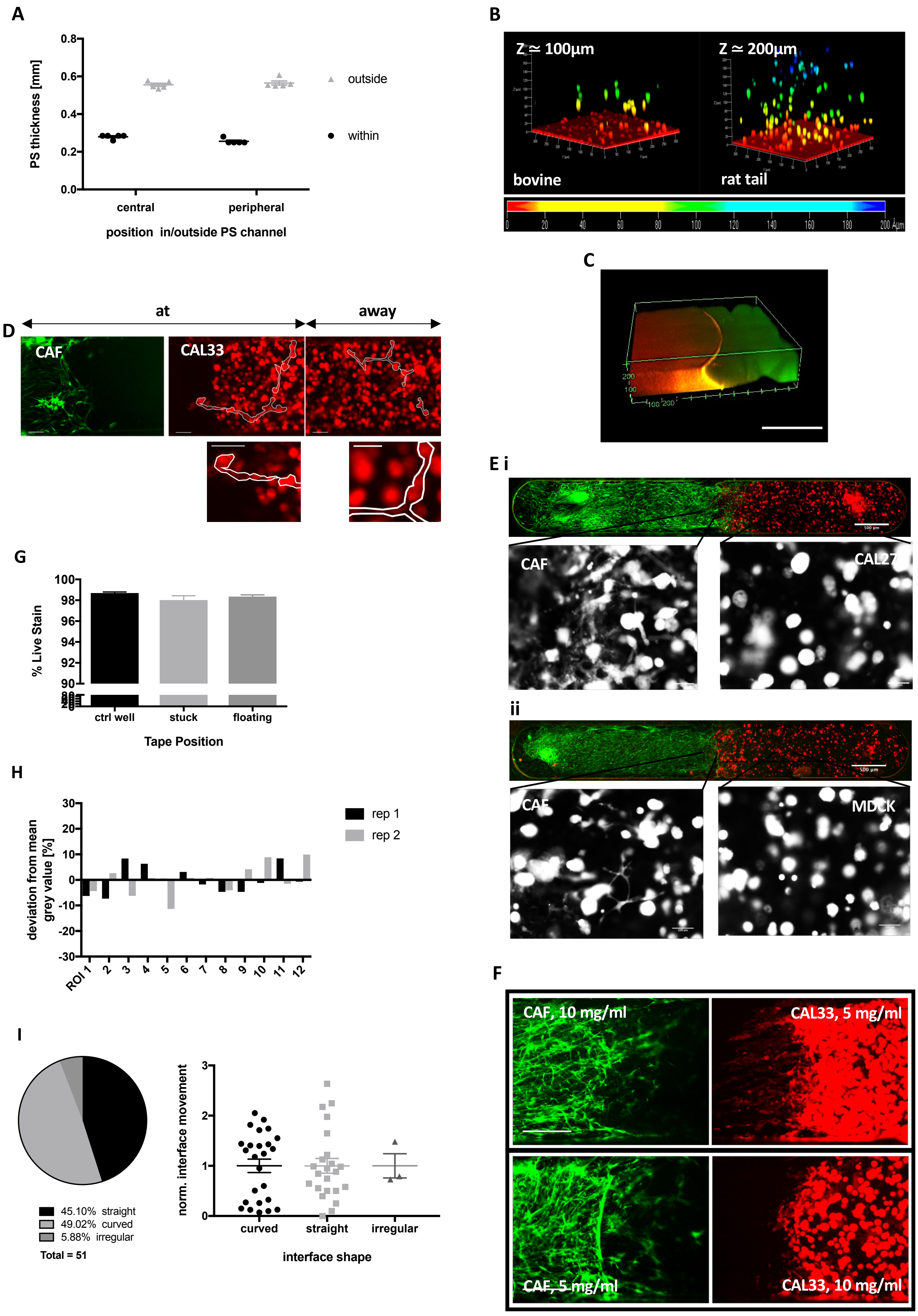
A
PS thickness [mm]
0.8
0.6
0.4
0.2
0.0
central
peripheral
position in/outside PS channel
outside
within
B
Z ≃ 100µm
Z ≃ 200µm
bovine
rat tail
0 20 40 60 80 100 120 140 160 180 200 µm
C
200
100
100 200
D
at
away
CAF
CAL33
E i
CAF
CAL27
ii
CAF
MDCK
500 µm
G
% Live Stain
100
98
96
94
92
90
ctrl well
stuck
floating
Tape Position
H
deviation from mean grey value [%]
30
20
10
0
-10
-20
-30
ROI 1 2 3 4 5 6 7 8 9 10 11 12
rep 1
rep 2
I
45.10% straight
49.02% curved
5.88% irregular
Total = 51
norm. interface movement
3
2
1
0
curved
straight
irregular
interface shape
F
CAF, 10 mg/ml
CAL33, 5 mg/ml
CAF, 5 mg/ml
CAL33, 10 mg/ml

**SI Figure 1 (**A) PS chip thickness consistency after hot embossing. Measurements were obtained using a caliper and measuring PS thickness inside and outside of PS open channels, at channel sites chosen randomly at the PS sheet center and periphery. Error bars represent SEM, n = 5 PS sheets. (B) Nuclear distribution as surrogate measure for cell distribution within bovine and rat tail collagen type 1 matrices. Images were taken directly after gelation of hydrogels of equal volume and cell density in PS open channels. Colder colours represent increasing distances from the PS channel bottom (red). (C) Hydrogel compartment continuity was assessed at the interface by fluorescently labelling both hydrogel compartments (green and red). No matrix gap could be identified and compartments were in intimate contact, as seen by the diffusion and mixing of fluorescent labels across the interface (yellow). Scale bar represents 500um. (D) Representative image of GLAnCE Matrigel culture. Tumor cell (red - mCherry) strand structures are outlined for clarity (white) and were present both, at the interface with CAFs (green - eGFP), and away. Scale bar represents 500 µm. Insets show strand structures in more detail. Scale bar is 250 µm. (E) GLAnCE compartmentalized culture generated with CAL27 cells and CAFs (i) and MDCK cells and CAFs (ii). Insets show cell morphology (white) at and away from the interface for CAL27 and MDCK cells. Scale bars represent 500 µm and 100 µm. (F) Representative interface images of GLAnCE hydrogels with dual stiffness. CAFs (green) were resuspended in 10 mg/ml (top panel) or 5 mg/ml rat tail type I collagen (bottom panel), with tumor cells (red) embedded in 5 mg/ml or 10 mg/ml collagen, respectively. Scale bar represents 500 µm. (G) Live-dead stain, using calcein AM and EthD-1, confirmed overall lack of cytotoxicity of the polyacrylic tape utilized for adhering the plate and the polystyrene chip, as assessed in 2D culture in standard 24 wells with and without tape added to the culture, either as a piece floating in the culture medium or stuck onto the well surface prior to cell seeding. Error bars represent SEM, n = 3 wells. (H) MTT signal from 2 GLAnCE wells was quantified as percentage deviation from the mean grey value measured in ROIs across the hydrogel length. Within-replicate consistency indicated homogeneous cell seeding and distribution at day 5 of culture. (I) Interface shapes were categorized within a representative subset of compartmentalized cocultures in GLAnCE, revealing 3 interface shapes, with straight and curved being predominant (pie chart). No correlation was found between interface shape and interface movement over time, indicating that all interfaces were suitable for analysis. Error represents SEM, total n= 51.

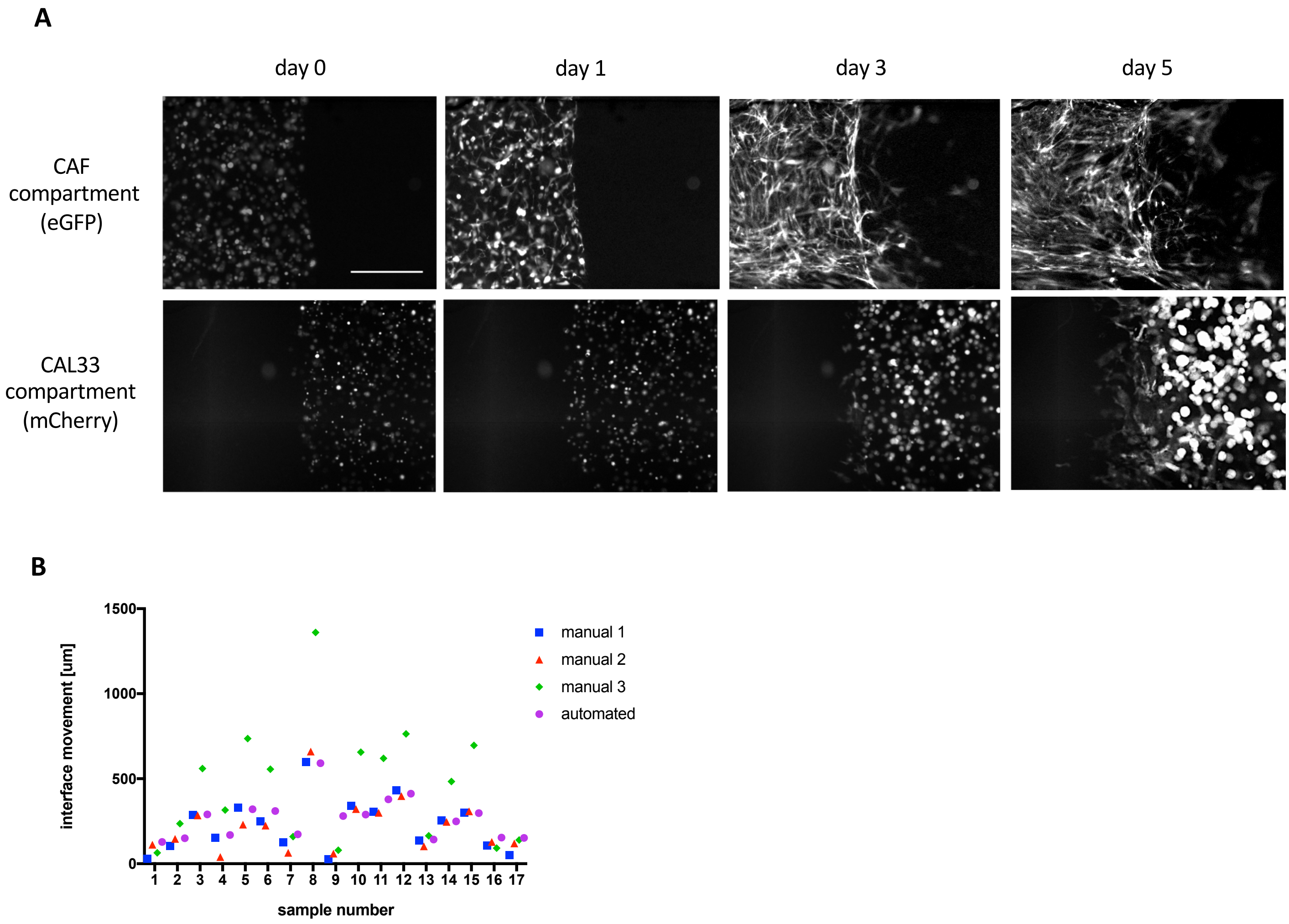


**SI Figure 2** (A) Time-lapse images of cell movement at the compartment interface over 5 days of coculture. The fluorescence image for each cell population is shown separately. Scale bar is 500 µm. (B) Comparison of automated and manual interface detection and interface movement quantification. Manual quantification was carried out on 17 random GLAnCE samples, by three users independently. Values from the automated quantification fell within the range and variability of the three manual calculation sets, showing an accuracy comparable to manual interface tracing.

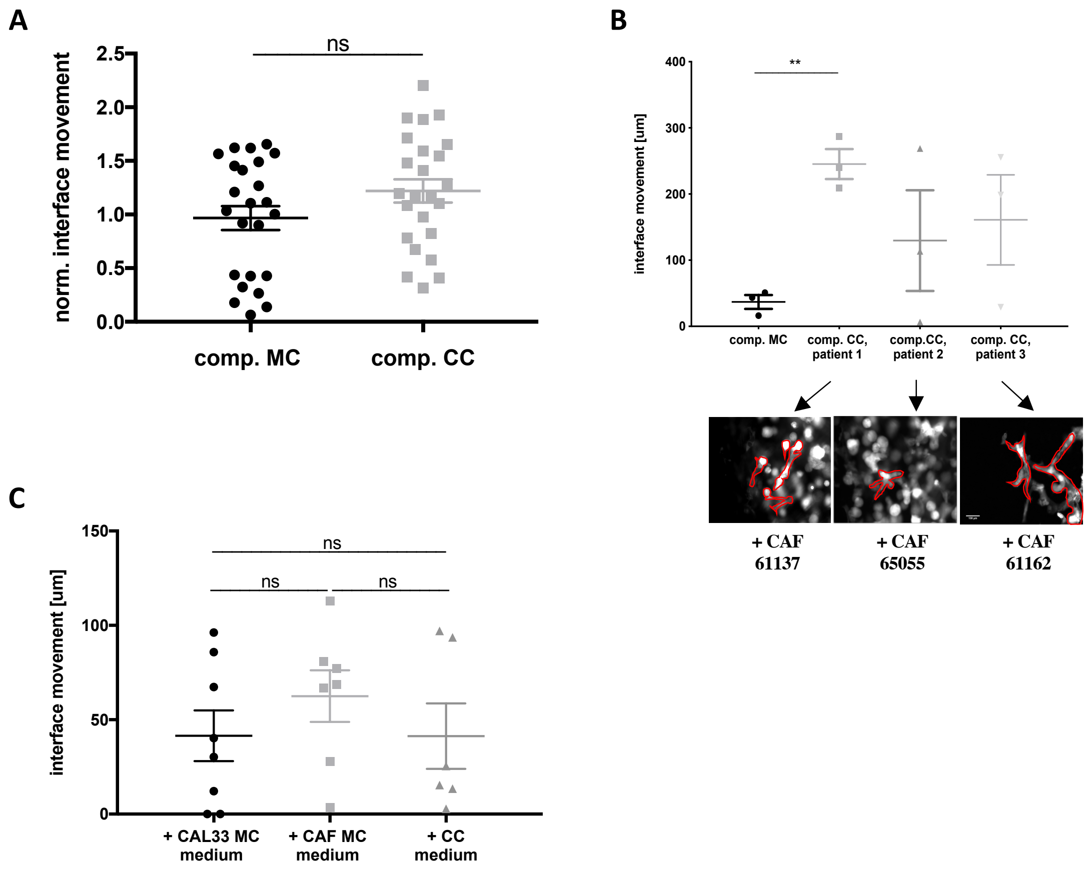


**SI Figure 3** (A) Inclusion of 2D, 'independent' clusters of tumor cells (defined as more than 50 pixels away from the interface; white in inset in Figure 2) into the analysis, obscured the differences in overall interface movement between tumor monoculture and coculture conditions. Error bars represent SEM, n=3. (B) Comparison of interface movement and invasive strand morphology of tumor cells cultured with 3 distinct, patient-derived CAF lines. While all CAFs lines elicited tumor invasiveness at the compartment interface (strands highlighted in red), only CAFs from patient 1 induced consistent interface expansion. Scale bar is 100 µm, error bars represent SEM, n = 3. (C) Conditioned media was harvested from tumor cell monocultures, CAF monocultures and cocultures, and was applied to compartmentalized tumor cell monocultures. None of the conditioned mediums led to a significant increase in interface movement compared to control.

**A**

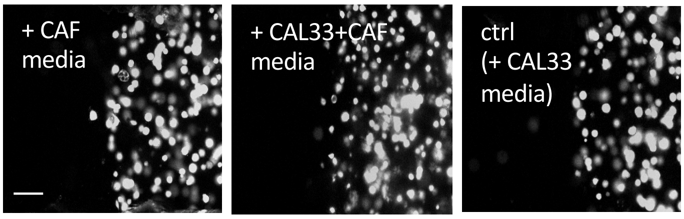


**SI Figure 4** Tumor (white) strand formation could not be induced in monocultures under treatment with conditioned media from CAFs nor CAF-tumor cell mixed cocultures, suggesting that CAF-secreted soluble factors were not responsible for invasive strand formation in compartmentalized cocultures. Scale bar represents 200 μm.

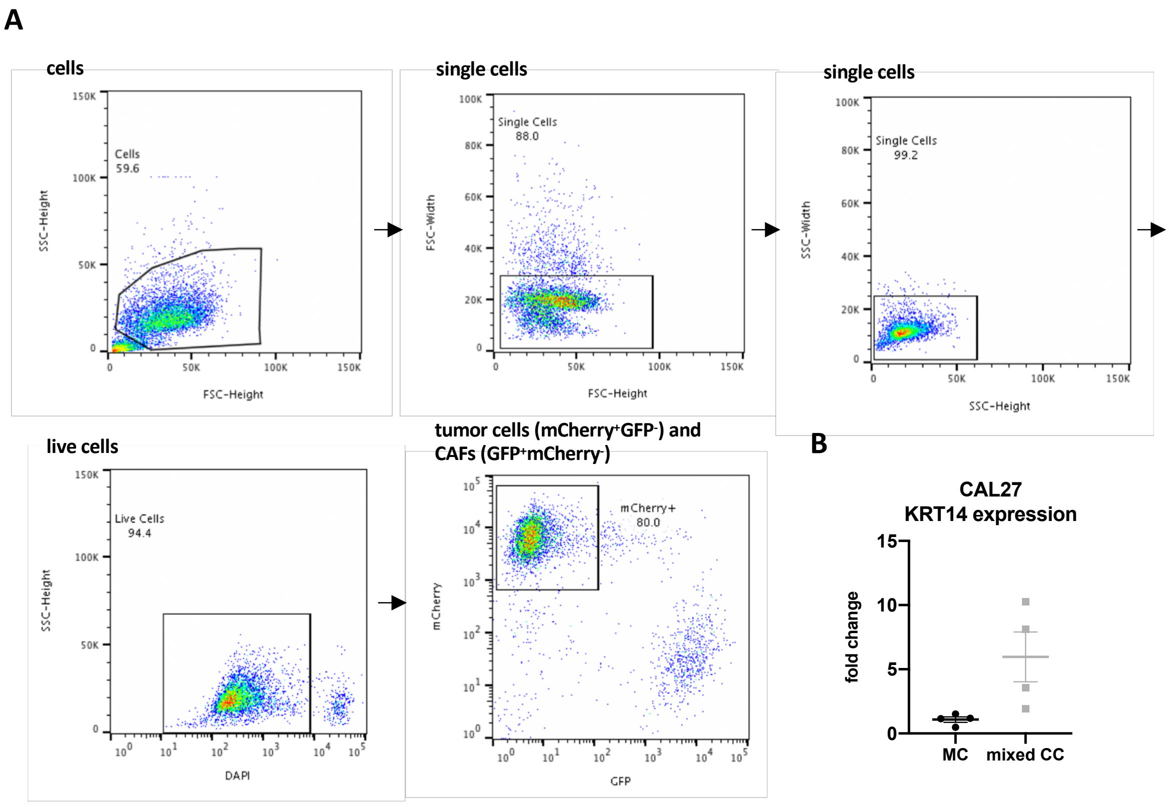


**SI Figure 5** (A) Gating path for isolating mCherry$^+$ tumor cells from tumor - CAF coculture GLAnCE samples. (B) Analysis of KRT14 in CAL27 HNSCC tumor cells shows increased (mean fold change >2) KRT14 expression in mixed CC, compared to tumor cell monocultures. Error bar is SEM, n=4.
